# Genetic variation in the odorant receptor gene OR4 and bit habits in natural populations of *Aedes aegypti* from Antioquia department, Colombia

**DOI:** 10.1101/552182

**Authors:** Juan Sebastian Pino

**Affiliations:** Molecular genetics laboratory, University of Antioquia

## Abstract

It has been determined factors that make humans more attractive to mosquitoes and which strategies they use to detect a potential host. Preferential differences for human/non-human hosts are related to variations in odorant receptors (OR) genes in the *Aedes aegypti* mosquitoes. This study use sequencing to establish the genetic variation in the odor receptor OR4 in 900 mosquitoes from different regions of Antioquia. A behavioral test using an olfactometer was also made to stablish the relationship of these variation with the attraction on different human hosts. The analysis in the attraction and OR4 variants did not show significant differences in the arrival rate among different human hosts. No significant differences in the description of OR4 variants between populations and between hosts, show that this gene is homogeneously distributed. The analysis showed a high genetic population diversity, measured as polymorphism and heterozygosity. This may be due to a few high frequency haplotypes in all the populations examined, suggesting a model of high gene flow between populations and/or selection in favor of these variants in all populations. Other low-frequency variants, many of which are population-specific, reflect the effect of genetic drift probably due to stochastic changes in the size of natural mosquito populations.

## Introduction

Mosquitoes belong to the most important group of disease-causing vectors. This is evidenced by the large number of species involved in the transmission of parasites and pathogens to humans and animals. Several of the most prevalent infectious diseases in the world, especially malaria, lymphatic filariasis and dengue, as well as less common diseases such as Japanese encephalitis, Chikungunya, Rift Valley fever, West Nile virus and the virus Usutu, are transmitted by them. Transmission between vertebrate hosts occurs through the blood feeding habit of mosquitoes, this allow that pathogens successfully established in a population and are transmitted by its arthropod guests. The habit of feeding on blood is part of its intrinsic character. Blood proteins are essential nutrients for egg production and reproductive efficiency (Chaves et al., 2010).

It has been reported that there are differences in the preference of mosquitoes for their hosts, resulting from a selective behavior, not only between different species, but also between populations of the same species and even within a population (Lyimo & Ferguson, 2009). Many mosquitoes that feed on blood do not show a specific preference for hosts, which suggests that the source of the blood and its quality are irrelevant, that is, they do not affect reproductive efficiency; however, other studies have shown that the quality of the blood, and therefore the origin of it, can affect reproductive performance (Lehane, 2005).

Blood serves as a source of metabolic energy, depending on the internal state of the insect (Takken et al., 1998). Some studies have shown that the quality of the blood and, therefore, the host species can affect the reproductive performance (Lehane, 2005). In addition, many of the disease agents transmitted by mosquitoes are host-specific (*Plasmodium falciparum, Plasmodium vivax, Wuchereria bancrofti*, Dengue virus). Therefore, host preference is more likely to be more common than previously assumed given the evolutionary association between the vector and the pathogen (Takken & Verhulst, 2013).

The behavior of *Aedes aegypti* has been extensively studied in the laboratory, but field studies of behavior with this mosquito are rare. For example, laboratory studies have shown a strong preference for *Ae. aegypti* for human odorants (Bernier et al., 2003; Geier et al., 1996), but the confirmation of this behavior from field studies is insufficient. Given the very important role of this species of mosquito in the transmission of dengue, detailed knowledge about the presumed association to the transmission of the disease by wild populations would be essential for the design of effective instruments for the control of mosquitoes.

The sense of smell is considered the most important aspect for the survival of mosquitoes and could be said to be related to biological effectiveness, having evolved to adapt to various lifestyles depending on the ecological niche (Cande et al., 2013; Hansson & Stensmyr, 2011; Nei et al., 2008). It is the efficiency of smell that makes blood-sucking mosquitoes, which feed on humans, specialists in the detection of their hosts and in the transmission of pathogens that cause disease. Mosquitoes detect odorants using several chemoreceptor families called ORs, which are the most studied in terms of function.

Recent studies have shown that the preference of mosquitoes for human blood is based on the abundance and sensitivity of a gene to detect odors in their antennae, which makes them more sensitive to the odor emitted by a person. An important study supports the theory that the evolution of the food preference in mosquitoes of the Aedes genus is due to a strong correlation with functional genetic variations in an odor receptor, the Or4 receptor, which has a high sensitivity for sulcatone, a component of the smell of the human body (Mcbride et all, 2014). However, no studies have been conducted in which genetic variants in this gene are evaluated and their possible association with the preference to feed in humans.

Understanding how variants in the OR4 gene can be related to *Ae. aegypti* to feed on humans can be of great importance because it would identify genetic clues in mosquitoes that are allowing the transmission of pathogens. For the receptor of odors gene OR4, it has been determined that it has 7 haplotypes (nomenclatures of A to G) and it was evidenced that 3 of them, the haplotypes A, B and G were in a higher frequency in mosquito populations that lived in urban areas and that they preferred to feed on humans (McBride et al., 2014), which shows that a differential expression of these alleles can be related to this behavior in the predilection to food.

In Colombia, Dengue has become an endemic disease in almost the entire territory. During the period between October 2009 and November 2010 the largest epidemic in the history of the country occurred, with a total of 157,202 cases of dengue, 221 confirmed deaths and a lethality of 2.26%, having a great impact on the health of the population (Protocol of surveillance in public health dengue (SIVIGILA, 2015).

According to the epidemiological reports of Sivigila, Antioquia is one of the departments in which the largest number of Dengue cases are reported annually. The incidence being variable among the subregions of the department; the last report of the National System of Surveillance in Public Health-SIVIGILA of 2015 shows the presence of Dengue in 83 municipalities of the Department and it is evident that the incidence among the subregions is lower in the Aburrá Valley (2.76%), the West (3.94%), Magdalena Medio (4.93) and Oriente (9.47%); and greater in Urabá (25.44%), Bajo Cauca (14.41%), Northeast (13.41%), Southwest (13.61%) and North (12.03%). In turn, within each subregion there is a variable incidence of the disease (Sivigilia, 2015).

Genetic variability is an important factor for the adaptation of an organism (Hiragi et al., 2009). Factors such as mutation, genetic drift, selection, migration and the reproduction system maintain genetic variation between and within the populations of a species (Souza, 2011). The study of the genetic diversity of the natural populations of the *Ae. aegypti* can provide a correlation of genetic characteristics with the vectorial potential of this species, helping in the acquisition of information for the development of strategic actions that reduce the transmission of dengue (Spenassatto, 2011).

At this time there are no studies that report whether the genetic variability in the OR4 receptor in natural populations of *Ae. aegypti* may be associated with reports of dengue cases in areas of high transmission, such as the department of Antioquia, where there are some municipalities reporting more cases than others. In addition, given the medical importance of this species of mosquito, establishing the relationship between genetic variation and variation in their dietary preferences on different host species and on different individuals of the same species, could contribute to the design of effective instruments for their control or to avoid contact with humans. Likewise, a preliminary step to propose to evaluate these associations is to know the genetic variation of this gene in natural populations of *Aedes aegypti.*

## Materials and methods

### Mosquitoes

Populations of *Ae. aegypti* mosquitoes used in this study were selected taking into account that they came from sites with differences in the reports of Dengue cases and in human density. The evaluated localities were San Rafael in the eastern subregion, with 12920 inhabitants and an incidence of 7.7% of dengue cases for 2015; Cisneros in the northeast, with 8998 inhabitants and an incidence of 11.1% of dengue cases; Anza in the west, with 7580 inhabitants and incidence of 13.2% of dengue cases; and Medellin in the Aburrá Valley, with 2457680 inhabitants and an incidence of 60.7% of dengue cases. In each site, 20 ovitraps were installed, distributed in a transect in the form of a zig-zag along the head of each municipality, with a distance of no more than 100 meters between each one. All the eggs collected in each population were transported to the insectary of the Medical Entomology Laboratory of the University of Antioquia; a batch of these eggs were hatched under standard insectary conditions (28°C temperature and 70% relative humidity). The adults emerged from each population were fed with mouse blood and crossed to obtain an F1 population with which all the tests were carried out.

### Analysis of the genetic variation of the olfactory receptor gene OR4 in populations

A sample of adult mosquitoes from each population was used to obtain DNA sequences from the OR4 gene region. For this, the individualized adult legs and thorax were taken to tubes labeled with a distinctive code, and from these the DNA was obtained with a modified salting-out protocol. Using standard protocols of molecular biology and carrying out modifications of the conditions described by McBride et al 2014, a pair of primers (AaegOR4-F5’GTTGACCTATTGCGTTTTCG3’; AaegOR4-R5’GCAAGTTCTGTTCTGATGTGC3’) were designed that amplify a region of 786 bp, which includes exons 2 and 3, with which we can distinguish the 7 haplotypes (A-G) described for this gene. The purifications of the amplicons and their nucleotide sequences were carried out by contracting a private service with Macrogen, Korea.

DNA sequences of the OR4 gene obtained were edited with Bioedit program v7.2.5 and polymorphic and heterozygous positions observed in the readings of the electropherograms were manually edited to give rise to a matrix of multilocus genotype data. As the gamification phase of the multilocus genotype data is unknown, the genotype data for each individual of each population were used to estimate the gametic phase by making a haplotype inference with the Bayesian ELB algorithm implemented in the Arlequin program v.3.5.1.2 described by Excoffier et.al (2003).

### Preferential analysis towards different human hosts

To evaluate the mosquito’s food source preference, an olfactometer Y shape was made, modified from the one originally described by Gouck, 1972 and the olfactometer made by Fernández-Grandon et al. (2015). The olfactometer was made of acrylic; consisting of a long tube (50 cm long and 7 cm internal diameter) attached to a release chamber (17 cm long) at the anterior end and to two Y-shaped arms at the posterior part. Each of the arms has a capture chamber (17 cm) and a test/control chamber (16 cm) in which the test odorants will be placed. The cameras in which the odorants are supplied, in this case a human hand, were attached to a constant flow of air that allowed the odor to spread throughout the olfactometer reaching the release chamber. The air was moistened in deionized water before being introduced into the two test branches of the olfactometer and its temperature was maintained at 25-27°C, with relative humidity of 60-80% and air velocity at 0.6 m s^-1^ (or 0.13 ms^-1^ in the arms and 0.11 ms^-1^ in the stem). This is considered the speed that encourages mosquitoes to fly against the wind.

Before conducting the tests, preliminary experiments were conducted to evaluate whether the mosquitoes chose each side (test chambers) of the olfactometer when they were given the same odor choice in both chambers (this in order to guarantee that both arms are functioning equally). It was also tested if mosquitoes could discriminate the side that contains an attractive odor when there is no odor on the other side. This in order to demonstrate that mosquitoes could follow the smell column of an attractant in the olfactometer.

To evaluate differences in the attraction to several human hosts, paired trials were conducted with six human male volunteers as baits, in each of the two compartments. Each of the six volunteers, who were selected with their ancestry in mind, participated in three paired trials, for a total of 15 trials (20 mosquitoes / trial). The volunteers were coded as H1 with African ancestry, H2 with European ancestry, H3 with Amerindian ancestry, H4 with African ancestry, H5 with European ancestry, and H6 with European ancestry. These 15 trials were replicated three times, for a total of 20 × 15 × 3 = 900 mosquitoes. In these experiments, mosquito entry rate in each of the two traps with odoriferous stimuli from two human volunteers was compared. Each volunteer was asked to avoid consuming alcohol, caffeine and fragrance products (perfume, deodorant, lotion, etc.) for 24 hours prior to testing. In addition, they were fully informed of the nature and purposes of the test.

For each test, batches of 20 adult females of *Ae. aegypti* of the F1 generation of the population of Medellin were used, 1-2 weeks after their emergence, not fed, without previous exposure to odors and previous contact with males to guarantee that the possible mating occurs; These were placed in a small cage before the trial to avoid post-handling stress. The females were fasted, without access to water for 16 to 24 hours before performing the tests. Ten minutes before the start of the trial, the mosquitoes were released in the main compartment to acclimate. At the end of the acclimation period, the air flow was ignited and a sliding door was opened between the capture chambers and main compartment to allow air to flow through the apparatus. The device allowed the mosquitoes to decide on the route to any of the odor stimuli, but they will not have physical contact with them.

With the door open, the mosquitoes could fly against the wind in response to the source of the attractant’s odor. At this point, they had the option of flying towards the test or to the control chamber. The number of mosquitoes in each trap was counted initially after 10 minutes; eventually this time was modified empirically to produce consistent responses over time. The stimuli alternated daily between the left and right ports to control the lateral bias.

The number of mosquitoes in each area of the olfactometer was recorded at the end of the tests. The following data were recorded: 1) the proportion of mosquitoes that flew against the wind and remained in the olfactometer stem; and 2) relative attraction (the proportion of mosquitoes in the arm that contained the odor stimulus).

### Genetic variation in the OR4 gene between different human hosts

The legs and thorax of a sample (10%) of individualized adults from each trap, in each of the 15 trials (H1-H2, H1-H3, H1-H4, H1-H5, H1-H6, H2-H3, H2-H4, H2-H5, H2-H6, H3-H4, H3-H5, H3-H6, H4-H5, H4-H6, H5-H6), were taken to tubes labeled with a distinctive code. From this sample, total DNA and OR4 sequences were obtained as described above. With these, the composition of haplotypes in each individualized host was estimated. The presence and frequency of OR4 haplotypes was used to obtain Fst estimates of genetic differentiation between individual hosts.

### Data analysis

The differences in the mosquito entry rate to each trap for each individual (and the average attraction rate) for each trial (differences between humans) were analyzed with the JMP package (SAS Institute Inc) through a t test. The data of the different replicas of the same test are presented as mean ± SE and a value of P <0.05 was considered statistically significant. For the statistical analysis of the preference, the significance of the difference between the test individuals was evaluated using a two-way t test, where the mean of each individual served as a single data point.

## Results

### Genetic diversity in the olfactory receptor gene OR4

The analyzed region showed a high diversity estimated as a level of polymorphism and heterozygosity. In a fragment of 525 bp in 175 total sequences, 54 polymorphic sites (10.28%) were observed, of which 26 showed some level of heterozygosis (Fig 1A, Table S1). A Hardy-Weinberg Equilibrium test showed that of those 26 heterozygous positions, 15 were in Hardy-Weinberg Imbalance in some population, mainly due to an excess of heterozygosis (Table S2). In the total sample, 107 haplotypes were observed: haplotype 1 corresponds to haplotypes A and B reported by McBride et al. 2014 (these haplotypes converge in this study because the region analyzed is smaller than the one reported in McBride et al 2014 and some polymorphisms that distinguish these two haplotypes were not included in this study). Only three haplotypes were found in high frequency (≥10 individuals, Hap1, Hap7 and Hap43); Hap1 (n = 43) was not found in Anzá; Hap43 (n = 13) was only observed only in Medellin and San Rafael and Hap 7 (n = 10) was not found in San Rafael. Table S1 details the information of the haplotypes found.

**Figure 1.**
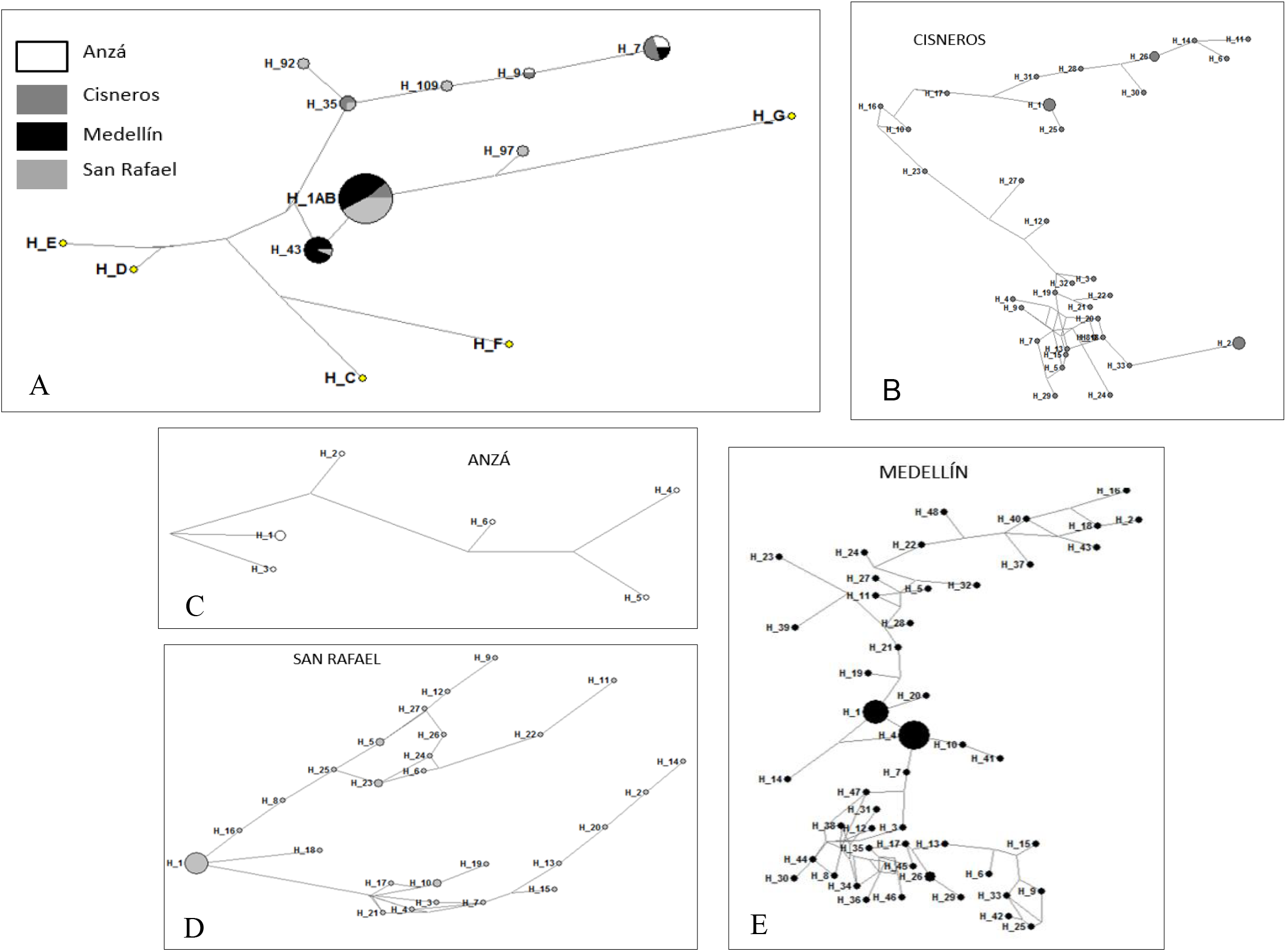
Haplotypes network of the 107 haplotypes found in the 175 sequences analyzed for the 4 populations studied. A) The most frequent haplotypes reported in this study, A-G haplotypes reported by McBride 2014 are included to show the relationship that exist between them and the haplotypes reported here, B) Most frequent haplotypes found in Cisneros municipality, C) Most frequent haplotypes found in Anza municipality, D) Most frequent haplotypes found in San Rafael municipality, E) Most frequent haplotypes found in Medellin municipality.

A network of more frequent haplotypes in the total sample showed that while the two most frequent haplotypes (Hap1 and Hap43) are separated by a mutational step, the Hap7 is phylogenetically distanced by several mutational steps (Fig 1A). Once the haplotype network was evaluated by comparing its frequency in the study populations, an intrapopulation analysis was carried out in order to evaluate the phylogenetic relationship of the most frequent haplotypes in each population (Fig 1B, C, D and E). In general, it is observed that haplotype 1 is the most frequent for each of the populations and that the rest of the haplotypes found are highly divergent, with some differences in the frequency of some of them; this is the case of the Hap2 and Hap26 in Cisneros, the Hap4 in Medellin and Hap5, Hap10 and Hap23 in San Rafael.

### Estimates of diversity of OR4 in the study populations

The estimates of diversity of these DNA sequences for each population are shown in Table 1. In Anzá population, the greatest genetic diversity was found, followed by Cisneros, Medellin and finally San Rafael (Table 1). The linkage disequilibrium test showed that 70 positions were in imbalance at a significance level of 0.05 after adjustment by Bonferroni (data not shown). The Fst estimate of genetic differentiation between populations based on both haplotype frequency and genotype composition showed significant differences between the populations of Anza/Medellin, Anza/San Rafael, Cisneros/Medellin and Cisneros/San Rafael (Table 2).

**Table 1.** Estimates of diversity in the sequences of the OR4 gene.

| Population | n | h | Hd | S | H | Pi | K | Hap. Esp. |
| --- | --- | --- | --- | --- | --- | --- | --- | --- |
| All | 175 | 107 | 0,93127 | 51 | 0.64 | 0.241 +/- 0.121 | 12,081 | NA |
| Anza | 8 | 6 | 0,89286 | 30 | 0.80 | 0.202 +/- 0.115 | 14,321 | 8 |
| Cisneros | 43 | 33 | **0,97453** | 39 | 0.71 | 0.200 +/- 0.101 | 13,671 | 43 |
| Medellin | 78 | 48 | 0,9207 | 47 | 0.69 | 0.169 +/- 0.086 | 11,559 | 78 |
| San Rafael | 46 | 27 | 0,8657 | 32 | 0.65 | 0.131 +/- 0.068 | 9,316 | 46 |

h: haplotypes; Hd: diversity of haplotypes; S: number of segregating sites; Pi: nucleotide diversity; H: heterozygosity; K: average number of differences

**Table 2.** Estimates of genetic differentiation among populations.

| Population | Fst |  |  |
| --- | --- | --- | --- |
|  | Frequency of haplotypes | Genotype | p |
| Anza-Cisneros | 0,035 | -0.00919 | 0.11712 |
| Anza-Medellin | **0,177** | **0.02622** | **0.00000** |
| Anza-San Rafael | **0,234** | **0.04307** | **0.00000** |
| Cisneros-Medellin | **0,083** | **0.01000** | **0.00000** |
| Cisneros-San Rafael | **0,092** | **0.01025** | **0.00000** |
| Medellin-San Rafael | 0,018 | -0.00186 | 0.03604 |

When translating these haplotypes to their corresponding reading frame, 38 protein haplotypes are produced. In these haplotypes there is a grouping between those that are more frequent and that are distributed in at least 3 of the 4 populations studied.

### Preference analysis between humans

The volunteers that participated in these experiments were fully informed of the nature and purposes of the test, emphasizing that at no time would they be in direct contact with the test mosquitoes. Table 4 shows the paired trials with each of the six volunteers, in each of the two compartments of the olfactometer. The mosquito entry rate was compared in each of the two traps of the olfactometer arms that had the odoriferous stimuli of two volunteers at the time. Before carrying out the tests with humans, all the standardization tests of the conditions inside the device were carried out, in order to avoid bias; these tests showed that when the mosquitoes undergo different stimuli they flew in an equal proportion towards the arms of the olfactometer and that when they only underwent one stimulus they flew in greater proportion towards that side of the apparatus. All these tests were done in triplicate, on different days and with different groups of mosquitoes.

**Table 3.**
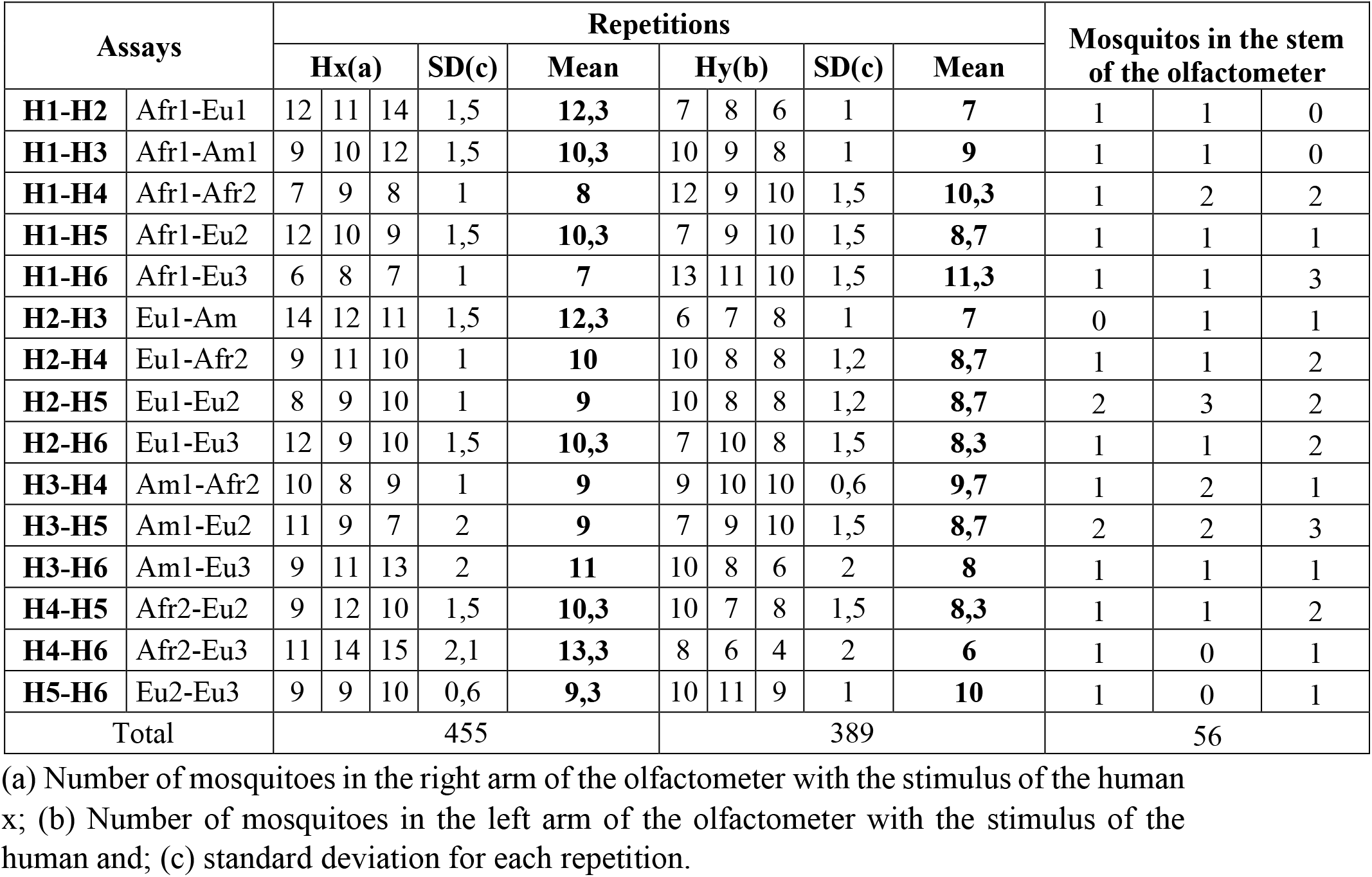
Relative attraction, represented by the number of mosquitoes flying towards each arm of the olfactometer in the three repetitions made for each paired test.

| Assays |  | Repetitions |  |  |  |  |  |  |  |  |  | Mosquitos in the stem of the olfactometer |  |  |
| --- | --- | --- | --- | --- | --- | --- | --- | --- | --- | --- | --- | --- | --- | --- |
|  |  | Hx(a) |  |  | SD(c) | Mean | Hy(b) |  |  | SD(c) | Mean |  |  |  |
| **H1-H2** | Afr1-Eu1 | 12 | 11 | 14 | 1,5 | **12,3** | 7 | 8 | 6 | 1 | **7** | 1 | 1 | 0 |
| **H1-H3** | Afr1-Am1 | 9 | 10 | 12 | 1,5 | **10,3** | 10 | 9 | 8 | 1 | **9** | 1 | 1 | 0 |
| **H1-H4** | Afr1-Afr2 | 7 | 9 | 8 | 1 | **8** | 12 | 9 | 10 | 1,5 | **10,3** | 1 | 2 | 2 |
| **H1-H5** | Afr1-Eu2 | 12 | 10 | 9 | 1,5 | **10,3** | 7 | 9 | 10 | 1,5 | **8,7** | 1 | 1 | 1 |
| **H1-H6** | Afr1-Eu3 | 6 | 8 | 7 | 1 | **7** | 13 | 11 | 10 | 1,5 | **11,3** | 1 | 1 | 3 |
| **H2-H3** | Eu1-Am | 14 | 12 | 11 | 1,5 | **12,3** | 6 | 7 | 8 | 1 | **7** | 0 | 1 | 1 |
| **H2-H4** | Eu1-Afr2 | 9 | 11 | 10 | 1 | **10** | 10 | 8 | 8 | 1,2 | **8,7** | 1 | 1 | 2 |
| **H2-H5** | Eu1-Eu2 | 8 | 9 | 10 | 1 | **9** | 10 | 8 | 8 | 1,2 | **8,7** | 2 | 3 | 2 |
| **H2-H6** | Eu1-Eu3 | 12 | 9 | 10 | 1,5 | **10,3** | 7 | 10 | 8 | 1,5 | **8,3** | 1 | 1 | 2 |
| **H3-H4** | Am1-Afr2 | 10 | 8 | 9 | 1 | **9** | 9 | 10 | 10 | 0,6 | **9,7** | 1 | 2 | 1 |
| **H3-H5** | Am1-Eu2 | 11 | 9 | 7 | 2 | **9** | 7 | 9 | 10 | 1,5 | **8,7** | 2 | 2 | 3 |
| **H3-H6** | Am1-Eu3 | 9 | 11 | 13 | 2 | **11** | 10 | 8 | 6 | 2 | **8** | 1 | 1 | 1 |
| **H4-H5** | Afr2-Eu2 | 9 | 12 | 10 | 1,5 | **10,3** | 10 | 7 | 8 | 1,5 | **8,3** | 1 | 1 | 2 |
| **H4-H6** | Afr2-Eu3 | 11 | 14 | 15 | 2,1 | **13,3** | 8 | 6 | 4 | 2 | **6** | 1 | 0 | 1 |
| **H5-H6** | Eu2-Eu3 | 9 | 9 | 10 | 0,6 | **9,3** | 10 | 11 | 9 | 1 | **10** | 1 | 0 | 1 |
| Total |  | 455 |  |  |  |  | 389 |  |  |  |  | 56 |  |  |

(a) Number of mosquitoes in the right arm of the olfactometer with the stimulus of the human x; (b) Number of mosquitoes in the left arm of the olfactometer with the stimulus of the human and; (c) standard deviation for each repetition.

**Table 4.** Estimates of diversity taking into account human volunteers.

| Host | n | h | Hd | S | Pi | K |
| --- | --- | --- | --- | --- | --- | --- |
| H1 (Afr1) | 16 | 9 | 0,8667 | 37 | 0,0211 | 11,075 |
| H2 (Eur1) | 13 | 8 | 0,85897 | 23 | 0,01324 | 6,9487 |
| H3 (Ame1) | 11 | 11 | **1** | 32 | 0,02403 | 12,618 |
| H4 (Afr2) | 13 | 11 | 0,96154 | 34 | 0,02786 | 14,641 |
| H5 (Eur2) | 11 | 10 | 0,98182 | 33 | 0,02192 | 11,509 |
| H6 (Eur3) | 14 | 10 | 0,9231 | 31 | 0,01924 | 10,099 |
| African | 29 | 17 | 0,894 +/- 0,043 | 46 | 0,02526 | 13,236 |
| European | 38 | 24 | 0,908 +/- 0,039 | 44 | 0,01882 | 9,881 |

h: haplotypes; Hd: diversity of haplotypes; S: number of segregating sites; Pi: nucleotide diversity; K: average number of differences

At the end of each trial, the number of mosquitoes in each area of the olfactometer was recorded and recorded the activity of flight against the wind, that is, the proportion of mosquitoes that flew against beyond 30 cm in the stem of the tube. This was reported as individuals who did not fly towards any of the stimuli provided (Table 4). These individuals at the time of delivery of this manuscript are in the process of resequencing because in a first shipment the samples could not be processed correctly.

The analyzes for the data obtained in the different tests were done with the JMP program through the t test for paired samples, this in order to establish if there were significant differences between the different volunteer hosts that were used in the experiments. When data was evaluated separately (t test of one tail) the p-value was significant (0.03), implying that mosquitoes have a tendency to prefer more to one host than to another; However, as this trend was evaluated in a paired analysis, the p-value (0.07) of the two-tailed test refutes this hypothesis, which is understood as that the mosquitoes evaluated do not have or show a marked predilection to choose a certain human host.

Taking into account the previous results and due to the availability of the sequences of the OR4 gene, tests were carried out to estimate the genetic diversity in the mosquitoes used for these experiments, for these tests the mosquitoes that preferred one human or another as the populations were considered. Table 5 shows the diversity estimates and it is observed that the individuals with the greatest diversity are Ame1, Eur2, followed by Afr2, Eur3 and finally Afr1 and Eur1.

**Table 5.**
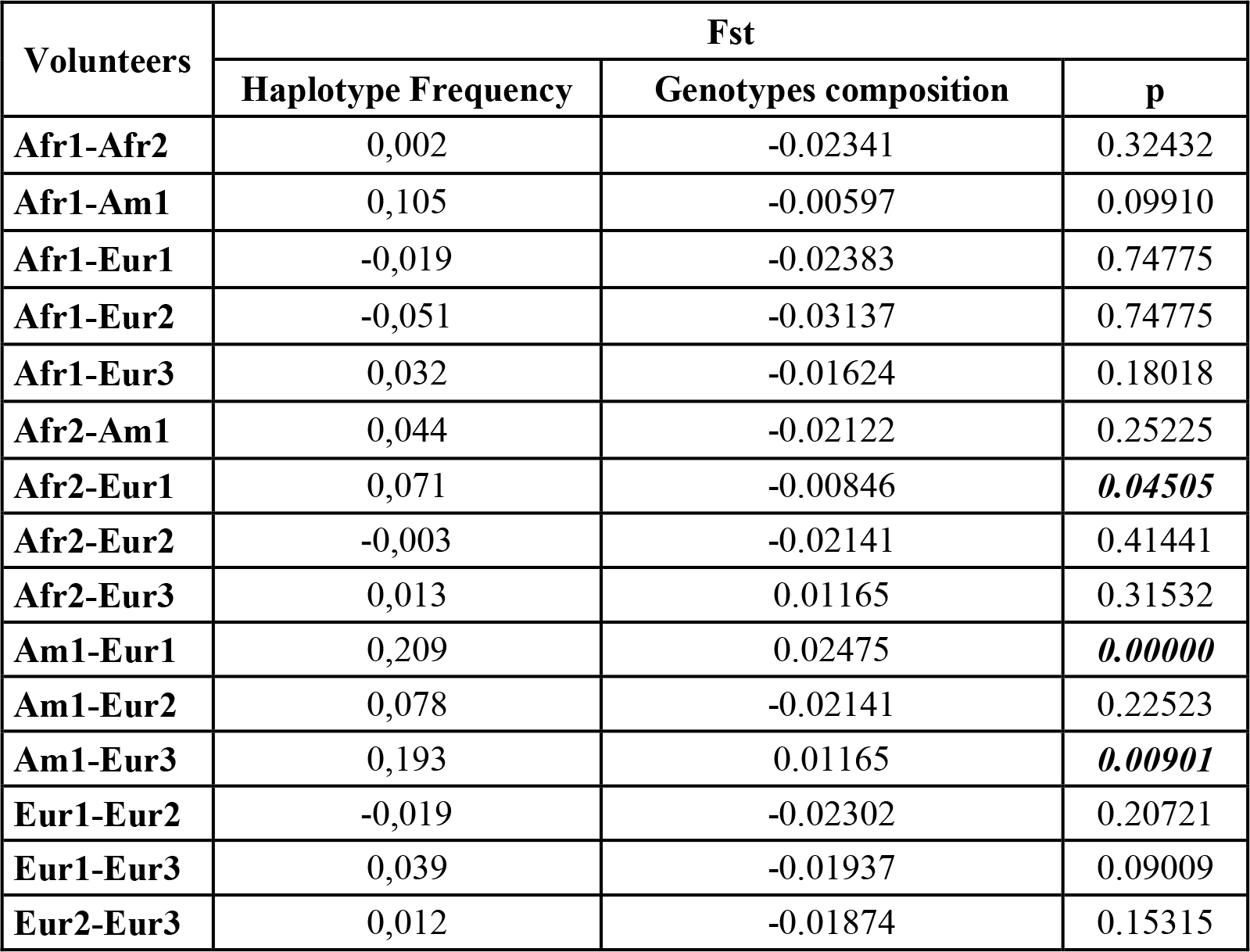
Estimates of genetic differentiation among individuals.

The analysis of genetic differentiation based on Fst estimates showed that both, haplotypes and OR4 genotypes differ significantly between individuals calling themselves as African1/European1 and Amerindian 1/European1 and European3 (Table 6); the other populations have low values of Fst, which is interpreted as there are no genetic differences between them and therefore there is greater homogeneity.

As with the haplotypes of the populations, those found in the preference analyzes were translated into their corresponding reading frame, producing 32 protein haplotypes. In these haplotypes there is a grouping among those that are more frequent and that is distributed in at least 2 of the 6 individuals studied.

## Discussion

In this study, a first analysis was made in order to establish the diversity of the OR4 gene in Anzá, Cisneros, Medellin and San Rafael populations; for this, the region of exons 2 and 3 of the OR4 gene was evaluated, which allows to establish the haplotypes reported by McBride et al (2014). It was found that for these populations there were 107 haplotypes, of which the most frequent haplotype was haplotype 1, which is related to haplotypes A and B of OR4, which have been associated with the preference of mosquitoes to feed on humans. Because the study populations were selected taking into account the reporting of dengue cases and that it was also expected to relate the frequency of haplotypes with this incidence, the result shows that in these populations are present the haplotypes that set a tendency to feed on humans, but there are too other factors involved in the transmission of the disease.

This haplotype 1 was not found in the population of Anzá, which may be due to the fact that in this population the mosquitoes present other haplotypes of the receptor or the small amount of sequences that were obtained once all the steps of filtering and editing were done, which was one of the limitations of this study. For the other two most frequent haplotypes, 43 was found only in Medellin and San Rafael and haplotype 7 was not found in San Rafael. In the case of Medellin this result coincides with the reporting of dengue cases, however, for San Rafael it is not. In this population it could be thought that although this haplotype is present, the transmission of the disease can be affected by other factors such as climate, which for this municipality is 22°C on average. The rest of haplotypes were found widely distributed in all populations and in low frequency.

The haplotype network in a geographical context showed that the most frequent haplotypes, 1 and 43, were separated by a mutational step, revealing the possible relationship between this haplotype 43 and the A and B haplotypes associated with human preference. When the haplotype network was evaluated by population, it was shown that some haplotypes increase their frequency, which suggests that some of them are specific for this population. This sense and taking into account the antecedent of the differential incidence of cases of dengue reported annually in the department of Antioquia, there is a possible association between these reports and the presence of the haplotypes of the OR4 gene. However, this it is an analysis that is worth performing at the intrapopulation level, increasing the number of mosquitoes evaluated in order to establish which are the surrounding haplotypes, the relationship between these haplotypes and to be able to pose experimentally how these may be conferring preference towards humans on mosquitoes.

This analysis allowed to demonstrate that this gene was highly polymorphic in the populations studied, and that some of its haplotypes, despite being in low frequency, are widely distributed in all populations. The high frequency of a few OR4 haplotypes in all populations and the low frequency of many population-specific haplotypes suggest expansions of the former and a possible genetic drift in the latter due to random geographic changes. Here we are talking about 4 populations from 4 municipalities of the Department, but it would be very interesting to be able to evaluate if this high level of polymorphism in the OR4 gene is maintained in other populations, taking into account the connectivity that exists between some regions and those where the Geographic barriers can facilitate the isolation of mosquitoes and therefore changes at the species level. The size of the analyzed sample does not allow to make exact conclusions about this effect.

54 polymorphic sites (10.28%) were observed, of which 26 showed some level of heterozygosis, which was evidenced in 15 sites in Hardy-Weinberg Imbalance in some population, mainly due to to an excess of heterozygosis. The polymorphic sites in this case may be those that are giving the identity to the large number of haplotypes found.

In general, it was observed that the populations of Anzá and Cisneros are differentiated from the populations of Medellin and San Rafael. This may be due to the geography of the department or the connectivity between these municipalities. In terms of epidemiology and anthropophilicity and in agreement, the frequency of the OR4 haplotypes found in this study could suggest that micro-geographic phenomena affecting transmission and preference are occurring in these populations.

The high diversity in Anzá is striking, even though it was the population with the lowest number of sequences. The Fst genetic differentiation analysis showed that among the Anzá and Cisneros populations, geographically more distant, they showed the lowest level of differentiation. In Colombia there are several works in which the population structure in *Ae. aegypti* (Atencia et al., 2018; Jaimes et al., 2015) taking into account mitochondrial sequences, it is in this sense that this work contributes to evidence that mosquito populations are also varying in genes that may have epidemiological importance, such as it is the OR4 which in the context of transmission of diseases such as Dengue, Zika or Chicungunya may be involved. It should be noted that due to its short flight range, the active dispersion of this mosquito is limited, especially between sites separated by great distances or between urban environments separated by wild environments, which could expect high genetic differentiation between distant populations. However, the passive dispersion fostered by the commercial connections of man can explain these unexpected patterns. Including connectivity among the municipalities of the department could also be understood better if this human dynamic is affecting the population dynamics of *Ae. aegypti*.

Even though the analyzes do not show marked differences between the haplotypes present, the similarity with the haplotypes reported in the literature as highly anthropophilic allow us to hypothesize that it would have an effect on the natural populations of mosquitoes, leading them to prefer feeding in humans and therefore increase the transmission of pathogens. In this sense, structural changes that occur between different haplotypes and how they interact with their respective odorant can be evaluated at the protein level; for this it is necessary to make a good bioinformatic approach and its subsequent experimental analysis.

It is often reported that when exposed to a group of hosts (of the same species), mosquitoes seeking hosts express a preference for one individual over others (Knols et al., 1995; Qiu et al., 2006; Takken & Knols, 2018). This preference is probably caused by the natural variation in odorants between individuals, which affects insects even at very low concentrations, and demonstrates the high sensitivity of odorant receptors to semiochemicals (Reisenman et al, 2016; Zwiebel & Takken,2004). However, the differences between species in the preference of mosquitoes by hosts are generally very robust because they are based on real differences in the composition of the odor (Majeed et al., 2016). In the tests of preference in this study it was found that for the human individuals used in the different paired analyzes there were no significant differences between the number of mosquitoes flying towards one host or another; this result could be due to the fact that variables that can modify the odor in humans were controlled and that the air flow with CO2 is not supplemented, this being an important variable as it has been determined as the main attraction for mosquitoes to detect a host.

The tests to estimate the genetic diversity in the mosquitoes used for these tests, considering the mosquitoes that preferred one human or another as the populations, showed that the individual with Amerindian ancestry presented the greatest diversity, followed by European 2 and African 2. This result supports what is mentioned in the previous paragraph, that is, probably the natural variation in the odorants is the one that is affecting the preference towards a host and that these function in interaction with the OR4 that must have a synergism with other receptors of this type; specifically, because individual odorants can activate specific groups of receptors while individual receptors can also respond to overlapping groups of odorants (Suh et al., 2014). Some receptors respond widely to a large number of odorants to act as “generalists”, while “specialized” ORs respond to sets of unitary or small odorants (Carey & Carlson, 2011). In addition, *Anopheles Gambiae* has reported ORs that are presumed to have evolved convergently, responding to odorants such as sulcatone and lactic acid (Carey et al., 2010), suggesting that the preference of the host is due to the selection of ORs profiles in the peripheral nervous system (Wolff & Jeffrey, 2018).

The analysis of genetic differentiation based on Fst estimates showed that both haplotypes and OR4 genotypes differ significantly between individuals calling themselves Africanno1 / European1 and Amerindian 1 / European 1 and 3. There are studies that have shown that body odor is affected by Ethnic differences in volatile organic compounds (VOC) among people of Caucasian, Oriental and Afro-American descent (Jacoby et al 2004; Martin et al 2010, Prokop-Prigge et al 2014, 2015). In this sense, it seems that ancestry is presenting an effect on the tendency of mosquitoes to look for humans; however, this possibility should be evaluated by linking an analysis with ancestry markers and mosquito preference trials.

The restriction of protein haplotypes towards certain ancestry could be evaluated in order to check if they are maintained and can change when the sample of mosquitoes, populations and human hosts evaluated increases. The analysis of the OR4 genetic variants among the mosquitoes used in the preference trials did not show significant differences in most inter-individual comparisons, however, when humans were evaluated as if they were affecting that choice it was observed that some individuals had differences among themselves; This may be due to the ancestry and odorants produced by each person, which vary in concentration and attractive effect.

Due to the limitations in the sample size for some populations, expanding the sample size and even the number of populations a broader analysis could be carried out to evaluate the composition of OR4 variants among populations, within the populations and between hosts. This is the first work in Colombia that evaluates or makes a first approach to the variability that mosquito odor receptors can present on the preference towards humans and how this could have an effect on the transmission of arboviruses. It is necessary to study the subject in order to find new strategies for public health prevention.

## Supplementary material

**Table S1.**
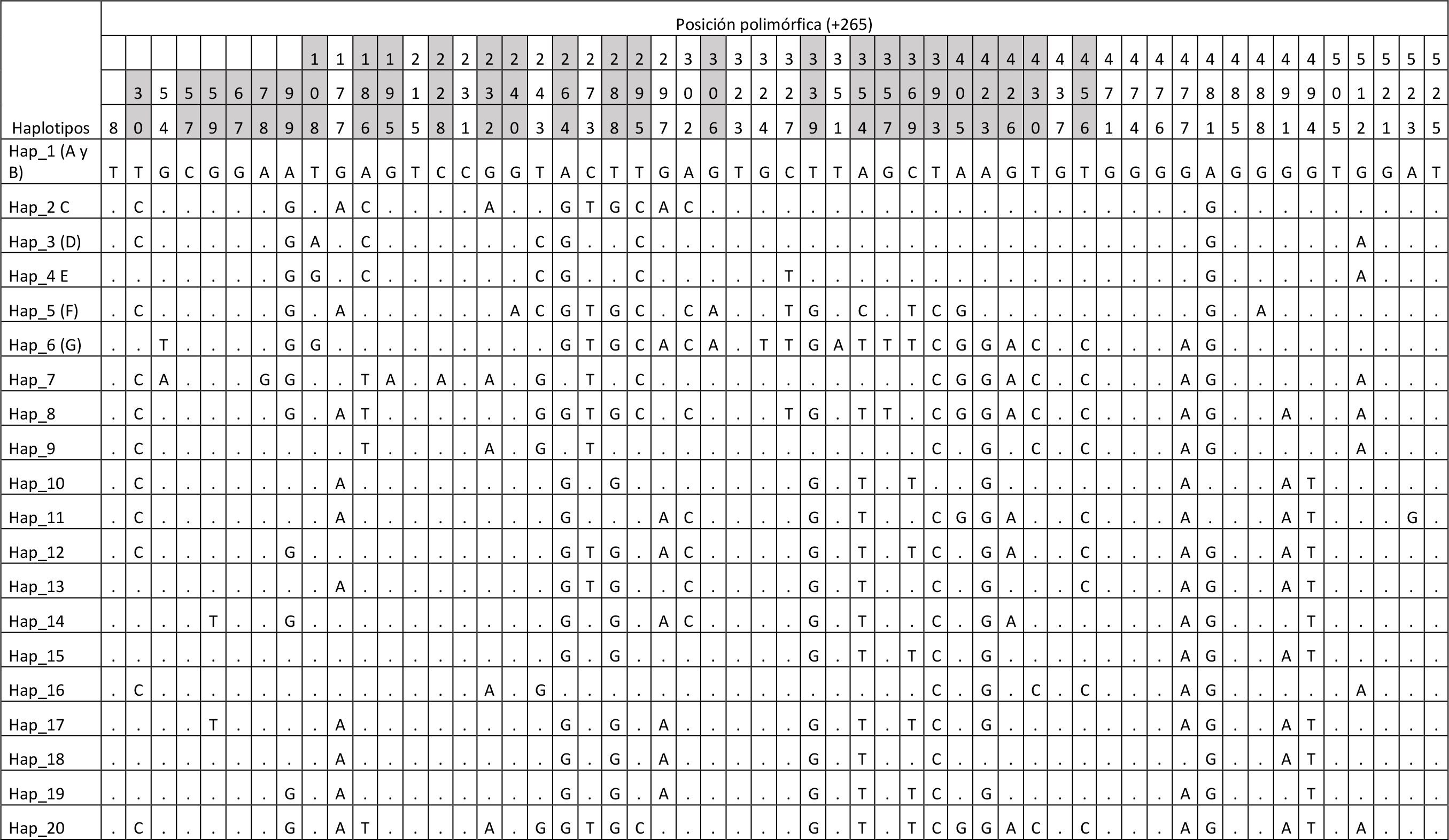
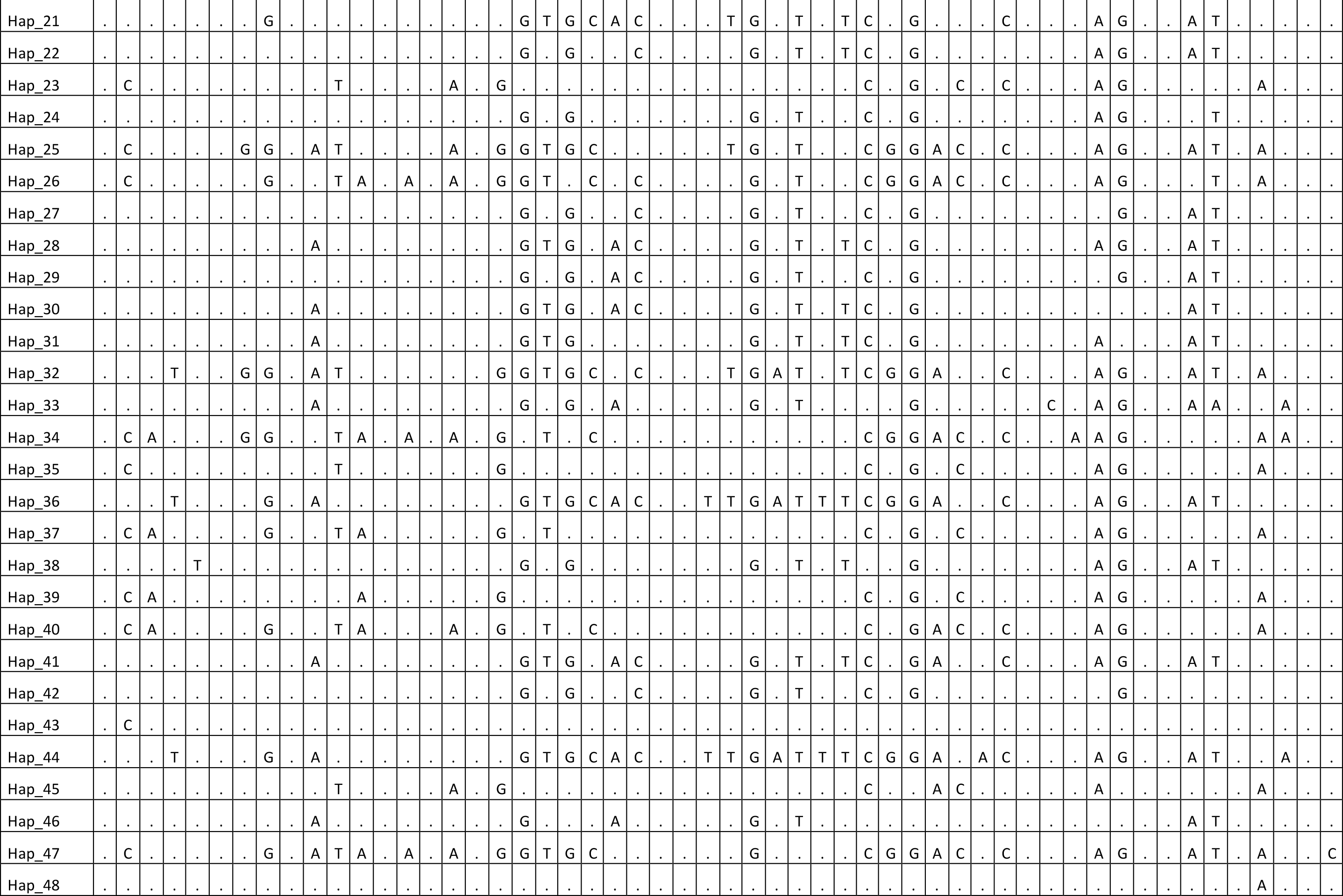
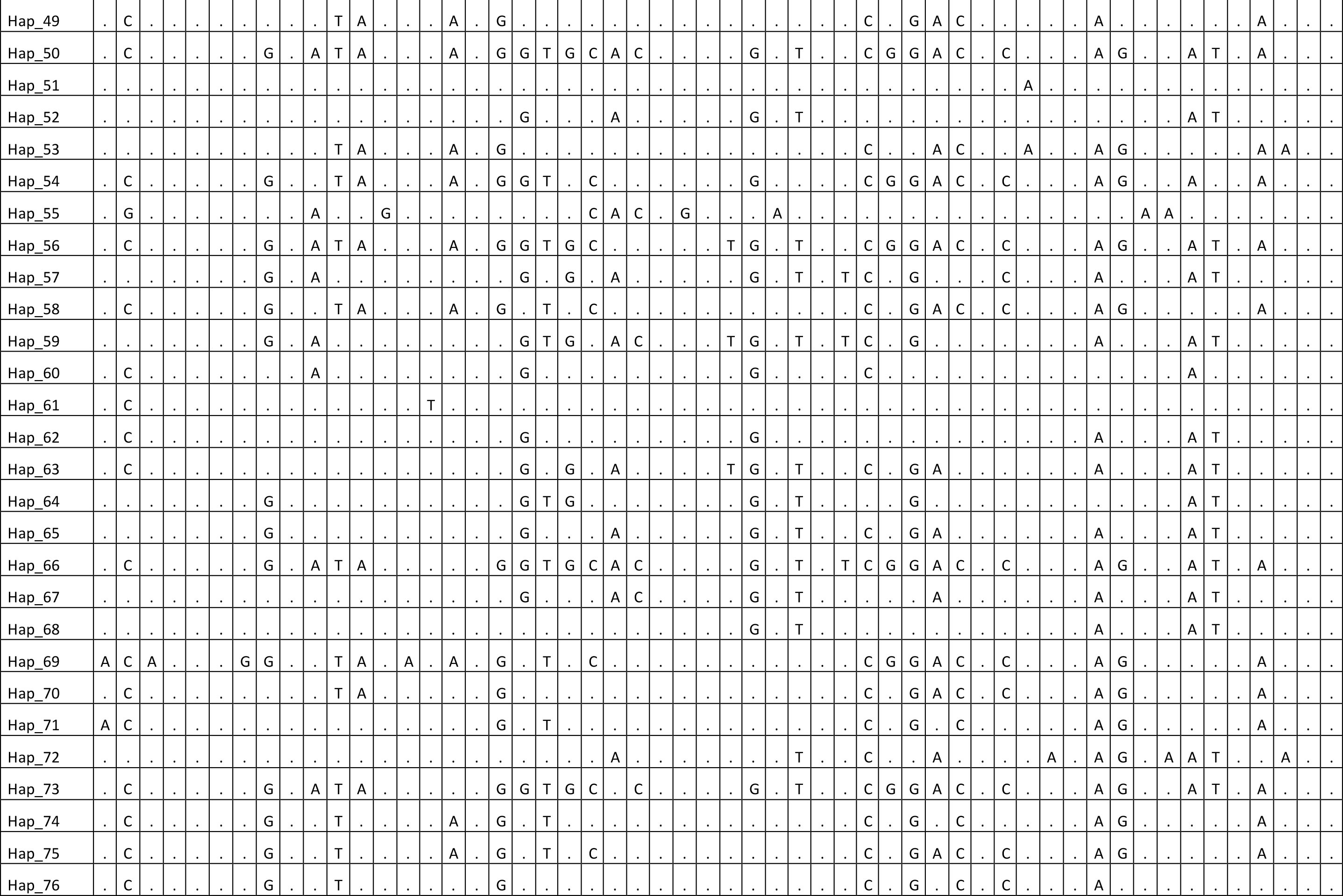
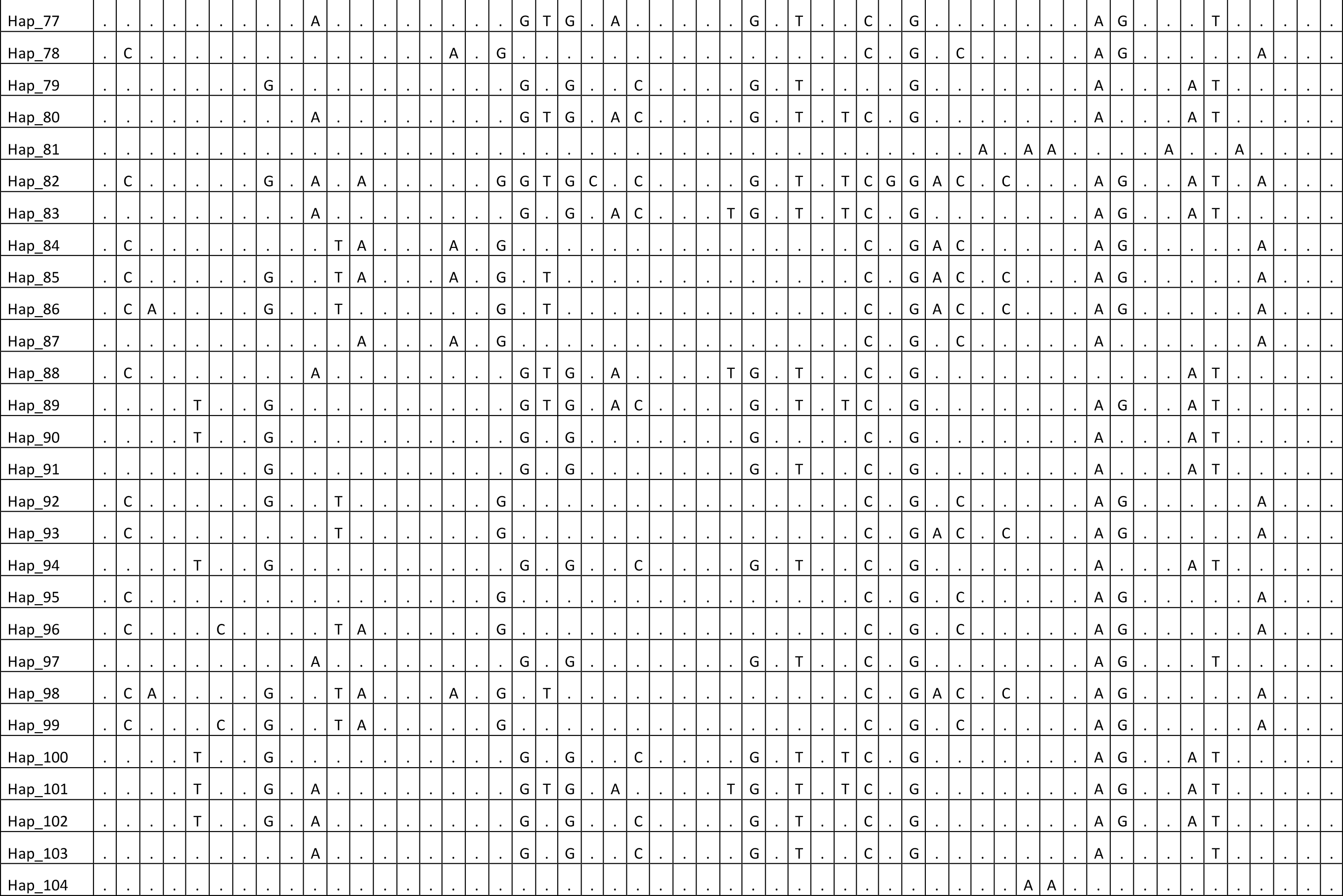
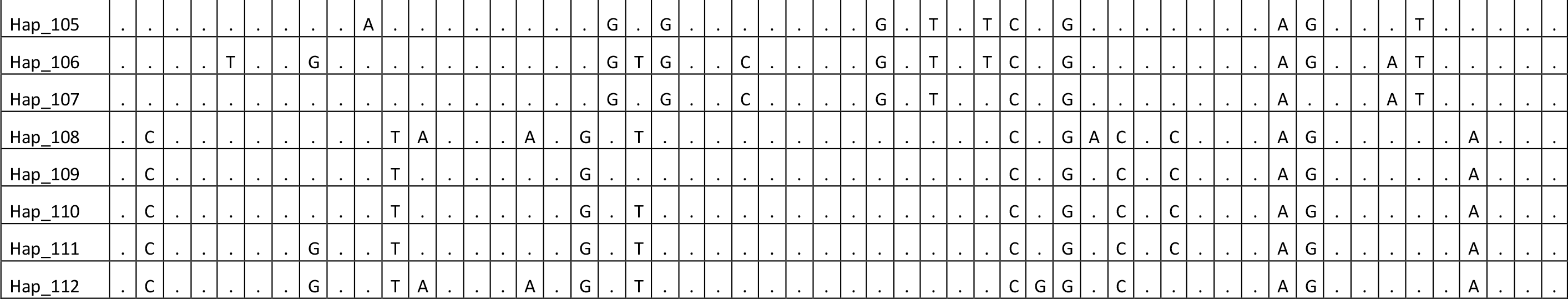
OR4 haplotypes. Polymorphic and heterozygous positions with respect to the reference sequences reported by McBride et al. 2014. The heterozygous positions are highlighted in gray.

**Table S2.**
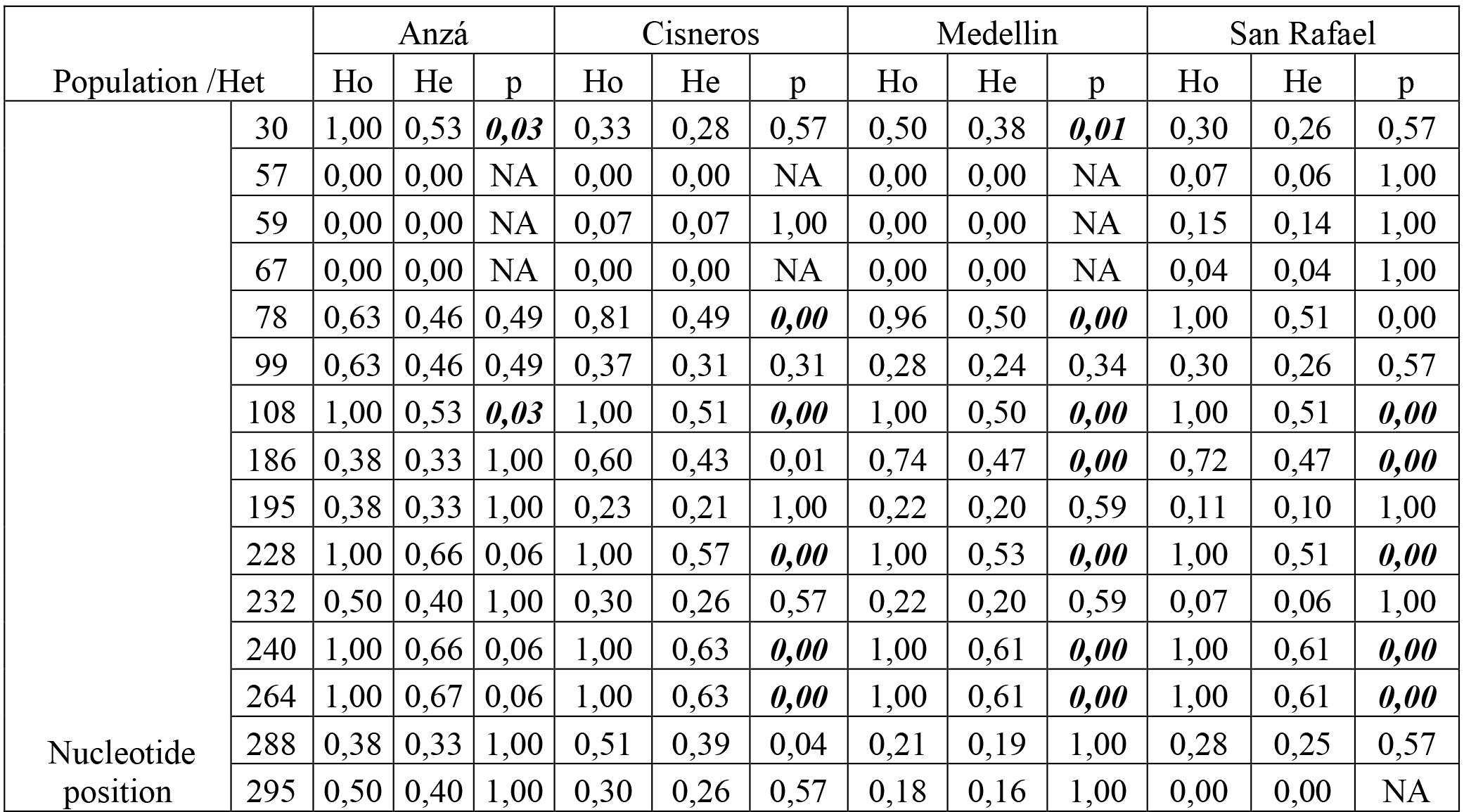
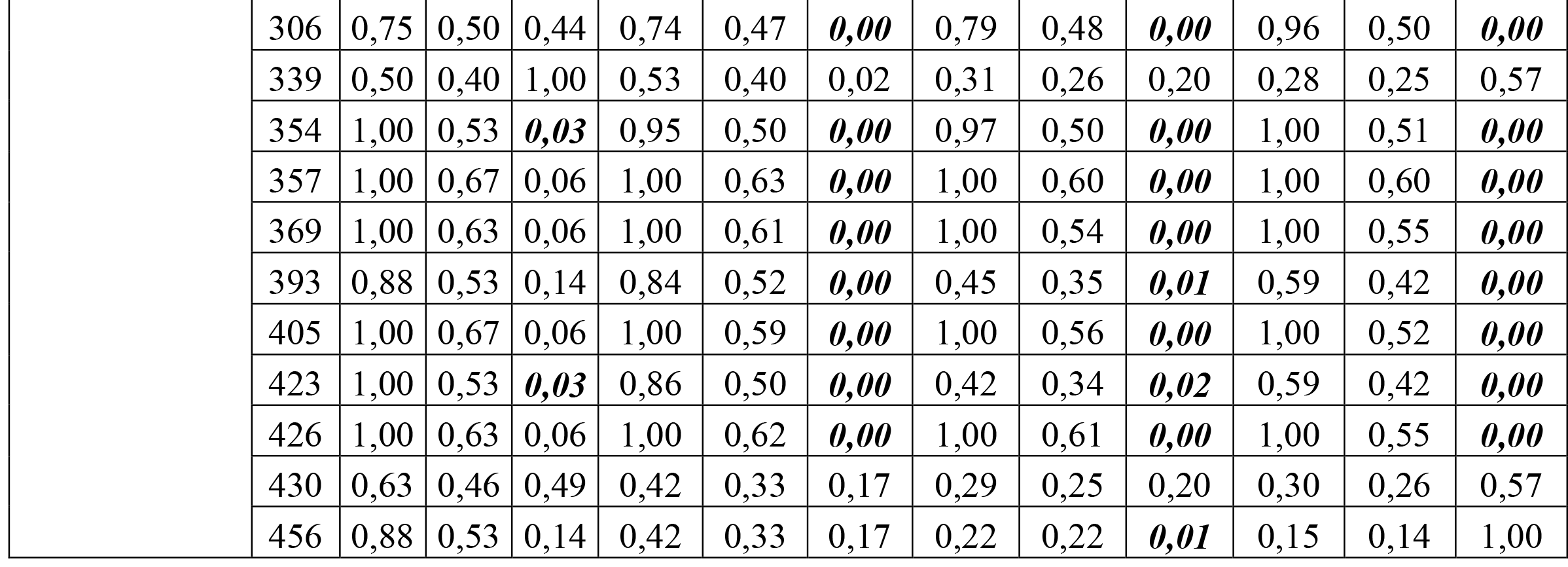
Heterozygosis and Hardy-Weinberg test.

| Population /Het |  | Anzá |  |  | Cisneros |  |  | Medellin |  |  | San Rafael |  |  |
| --- | --- | --- | --- | --- | --- | --- | --- | --- | --- | --- | --- | --- | --- |
|  |  | Ho | He | p | Ho | He | p | Ho | He | p | Ho | He | p |
| Nucleotide position | 30 | 1,00 | 0,53 | **0,03** | 0,33 | 0,28 | 0,57 | 0,50 | 0,38 | **0,01** | 0,30 | 0,26 | 0,57 |
|  | 57 | 0,00 | 0,00 | NA | 0,00 | 0,00 | NA | 0,00 | 0,00 | NA | 0,07 | 0,06 | 1,00 |
|  | 59 | 0,00 | 0,00 | NA | 0,07 | 0,07 | 1,00 | 0,00 | 0,00 | NA | 0,15 | 0,14 | 1,00 |
|  | 67 | 0,00 | 0,00 | NA | 0,00 | 0,00 | NA | 0,00 | 0,00 | NA | 0,04 | 0,04 | 1,00 |
|  | 78 | 0,63 | 0,46 | 0,49 | 0,81 | 0,49 | **0,00** | 0,96 | 0,50 | **0,00** | 1,00 | 0,51 | 0,00 |
|  | 99 | 0,63 | 0,46 | 0,49 | 0,37 | 0,31 | 0,31 | 0,28 | 0,24 | 0,34 | 0,30 | 0,26 | 0,57 |
|  | 108 | 1,00 | 0,53 | **0,03** | 1,00 | 0,51 | **0,00** | 1,00 | 0,50 | **0,00** | 1,00 | 0,51 | **0,00** |
|  | 186 | 0,38 | 0,33 | 1,00 | 0,60 | 0,43 | 0,01 | 0,74 | 0,47 | **0,00** | 0,72 | 0,47 | **0,00** |
|  | 195 | 0,38 | 0,33 | 1,00 | 0,23 | 0,21 | 1,00 | 0,22 | 0,20 | 0,59 | 0,11 | 0,10 | 1,00 |
|  | 228 | 1,00 | 0,66 | 0,06 | 1,00 | 0,57 | **0,00** | 1,00 | 0,53 | **0,00** | 1,00 | 0,51 | **0,00** |
|  | 232 | 0,50 | 0,40 | 1,00 | 0,30 | 0,26 | 0,57 | 0,22 | 0,20 | 0,59 | 0,07 | 0,06 | 1,00 |
|  | 240 | 1,00 | 0,66 | 0,06 | 1,00 | 0,63 | **0,00** | 1,00 | 0,61 | **0,00** | 1,00 | 0,61 | **0,00** |
|  | 264 | 1,00 | 0,67 | 0,06 | 1,00 | 0,63 | **0,00** | 1,00 | 0,61 | **0,00** | 1,00 | 0,61 | **0,00** |
|  | 288 | 0,38 | 0,33 | 1,00 | 0,51 | 0,39 | 0,04 | 0,21 | 0,19 | 1,00 | 0,28 | 0,25 | 0,57 |
|  | 295 | 0,50 | 0,40 | 1,00 | 0,30 | 0,26 | 0,57 | 0,18 | 0,16 | 1,00 | 0,00 | 0,00 | NA |

|  |  |  |  |  |  |  |  |  |  |  |  |  |
| --- | --- | --- | --- | --- | --- | --- | --- | --- | --- | --- | --- | --- |
| 306 | 0,75 | 0,50 | 0,44 | 0,74 | 0,47 | **0,00** | 0,79 | 0,48 | **0,00** | 0,96 | 0,50 | **0,00** |
| 339 | 0,50 | 0,40 | 1,00 | 0,53 | 0,40 | 0,02 | 0,31 | 0,26 | 0,20 | 0,28 | 0,25 | 0,57 |
| 354 | 1,00 | 0,53 | **0,03** | 0,95 | 0,50 | **0,00** | 0,97 | 0,50 | **0,00** | 1,00 | 0,51 | **0,00** |
| 357 | 1,00 | 0,67 | 0,06 | 1,00 | 0,63 | **0,00** | 1,00 | 0,60 | **0,00** | 1,00 | 0,60 | **0,00** |
| 369 | 1,00 | 0,63 | 0,06 | 1,00 | 0,61 | **0,00** | 1,00 | 0,54 | **0,00** | 1,00 | 0,55 | **0,00** |
| 393 | 0,88 | 0,53 | 0,14 | 0,84 | 0,52 | **0,00** | 0,45 | 0,35 | **0,01** | 0,59 | 0,42 | **0,00** |
| 405 | 1,00 | 0,67 | 0,06 | 1,00 | 0,59 | **0,00** | 1,00 | 0,56 | **0,00** | 1,00 | 0,52 | **0,00** |
| 423 | 1,00 | 0,53 | **0,03** | 0,86 | 0,50 | **0,00** | 0,42 | 0,34 | **0,02** | 0,59 | 0,42 | **0,00** |
| 426 | 1,00 | 0,63 | 0,06 | 1,00 | 0,62 | **0,00** | 1,00 | 0,61 | **0,00** | 1,00 | 0,55 | **0,00** |
| 430 | 0,63 | 0,46 | 0,49 | 0,42 | 0,33 | 0,17 | 0,29 | 0,25 | 0,20 | 0,30 | 0,26 | 0,57 |
| 456 | 0,88 | 0,53 | 0,14 | 0,42 | 0,33 | 0,17 | 0,22 | 0,22 | **0,01** | 0,15 | 0,14 | 1,00 |

